# EVfold.org: Evolutionary Couplings and Protein 3D Structure Prediction

**DOI:** 10.1101/021022

**Authors:** Robert Sheridan, Robert J. Fieldhouse, Sikander Hayat, Yichao Sun, Yevgeniy Antipin, Li Yang, Thomas Hopf, Debora S. Marks, Chris Sander

## Abstract

Recently developed maximum entropy methods infer evolutionary constraints on protein function and structure from the millions of protein sequences available in genomic databases. The EVfold web server (at EVfold.org) makes these methods available to predict functional and structural interactions in proteins. The key algorithmic development has been to disentangle direct and indirect residue-residue correlations in large multiple sequence alignments and derive direct residue-residue evolutionary couplings (EVcouplings or ECs). For proteins of unknown structure, distance constraints obtained from evolutionarily couplings between residue pairs are used to *de novo* predict all-atom 3D structures, often to good accuracy. Given sufficient sequence information in a protein family, this is a major advance toward solving the problem of computing the native 3D fold of proteins from sequence information alone.

**Availability:** EVfold server at http://evfold.org/

**Abbreviations:**

DI: direct information
EC: evolutionary coupling
EV: evolutionary
MSA: multiple sequence alignment
PLM: pseudo-likelihood maximization
PPV: positive predictive value (number of true positives divided by the sum of true and false positives)
TM-score: template modeling score

## 1 INTRODUCTION

Evolution of species is now revealed at the molecular level through genomic sequencing. Extensive output from high-throughput sequencing has made it increasingly productive to mine this data for interactions affecting protein structure and function. Knowledge of which protein residues are involved in functionally important interactions and how these are arranged in space benefits research in many areas of biology. A long-standing goal in computational biology has been to predict 3D structure from amino acid sequence alone. As the Critical Assessment of Techniques for Protein Structure Prediction (CASP) has shown, accurate structure prediction remains a largely unsolved challenge, especially in the absence of a homologous template [1]. In addition, predicting the identity and role of functional residues in incompletely characterized proteins is another persistent challenge [2]. However, recent developments have led to a breakthrough in computational methods employing co-evolution [3–6] for both structure and function prediction. The use of evolutionary couplings (ECs) between residues to accurately predict all-atom protein structures was, to our knowledge, first demonstrated in the fall of 2010 and published in 2011 [7].

The EVcouplings method uses a global probability model for an isostructural set of protein sequences in the form of an exponential model, which has a pseudo-energy expression up to second order (residue pair interactions) as the exponent. The parameters in the model are extracted (not fit in the sense of machine learning) from co-variation counts for all pairs of residue positions in a multiple sequence alignment (MSA) of evolutionarily related protein sequences. The inferred ECs disambiguate direct from indirect correlations. Importantly, ECs between residue pairs are often involved in key functional and structural interactions that in general cannot be detected using single-column conservation within an MSA. ECs often occur between residues in structural contact, enabling their use in *de novo* predictions of protein structure. Complementary to the EVold server, which commenced public operation in April 2013, there are now several additional online resources for protein contact prediction [8–10]. Development of methods and applications is very active in the field of evolutionary couplings, including focused efforts on beta-barrel membrane proteins [11], protein complexes [12, 13] and hybrid methods for structure determination, such as combining sparse NMR data with residue-residue ECs (EC-NMR) [14].

## 2 EVOLUTIONARY COUPLINGS AND PROTEIN 3D STRUCTURES

### 2.1 Capabilities: structures and functional interactions

For a target protein sequence, embedded in a protein family alignment, the EVfold server derives ECs, which reflect structural and functional constraints. From the residue pair constraints, the server can *de novo* model 3D structures from sequence information alone (Figure 1), without the need for the 3D structure of homologous proteins or protein fragments, as in template model building or model building by homology.

**Figure 1:**
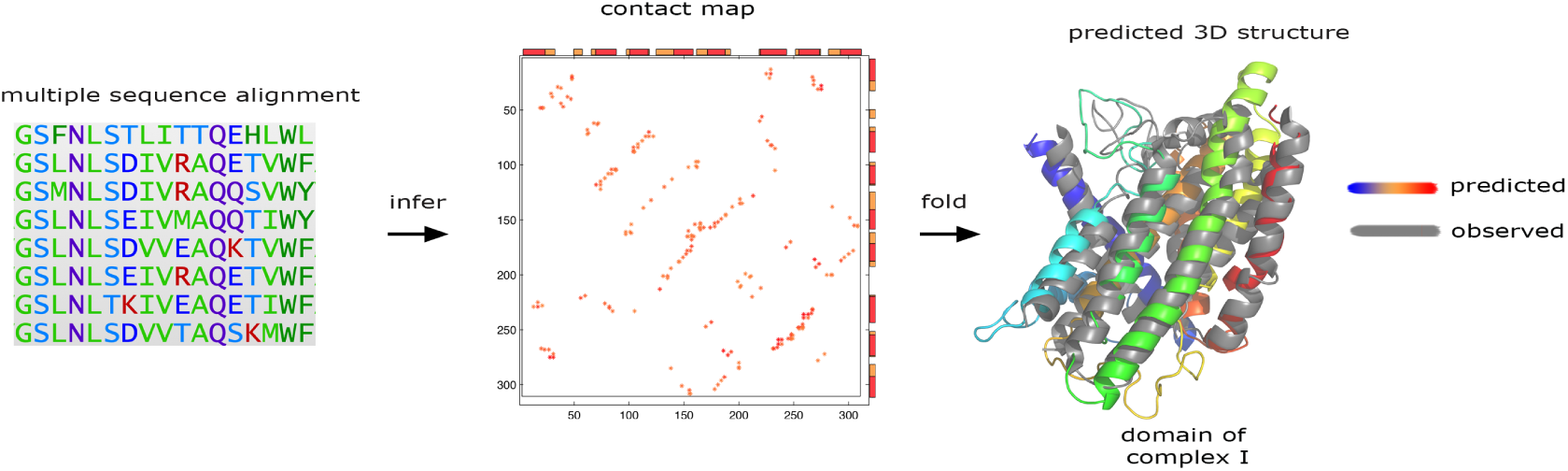
EVfold server process: from amino acid sequences to all-atom 3D structures (option EVfold). User supplies a sequence of interest (the target sequence) in the context of a large multiple sequence alignment (left) that provides residue-residue covariation information. Evolutionary constraints (ECs) are computed using DI or PLM (see Method) and the top ECs are predictive of residue-residue contacts (middle). The predicted 3D fold of the human subunit one of Complex I (right, gene name: MT-ND1, Uniprot: NU1M_HUMAN) [26] agrees well with the subsequently published crystal structure of the *Thermus thermophilus* homolog (Uniprot: NQO8_THET8, PDB: 4HE8) [27].

Beyond their use in computing 3D structures, the inferred ECs can be used to identify functionally constrained residue interactions indicative of active sites, protein-protein interfaces and other functional sites, and can be used to guide or interpret experiments that measure the phenotypic consequences of residue substitutions. The server can handle 3D structure prediction for globular and helical transmembrane proteins, with server capability for beta-barrel membrane proteins and for protein complexes technically feasible [11, 12], but not yet implemented.

### 2.2 Input: sequences

Minimal server input is a specific protein (database ID or amino acid sequence) and a sequence range (domain) within the protein. Computing ECs for a target protein sequence requires a MSA for a set of proteins plausibly isostructural with the target protein, typically paralogs from the same organism and homologs from other species. The MSA can be provided by the user or is retrieved from the Pfam domain database [15], or generated using software that searches for homologs in protein sequence databases, such as HHblits [16] or jackhmmer [17]. The depth (number and diversity of sequences) and breadth (coverage of the target protein domain by the aligned residues) of the protein family alignment has to be sufficient to allow reliable extraction of ECs. Typically, we suggest at least 5L sequences in the MSA, where L is the number of residues of the target protein domain and 75% breadth of coverage. For example, we suggest a 200 residue target protein have at least 1000 sequences in the MSA. For unknown structures, the server automatically predicts secondary structure using PSIPRED [18] and alpha-helical transmembrane topology using MEMSAT-SVM [19] and uses these as input to the 3D structure generation. Users can override the secondary structure predictions using experimental data or other predictions.

If an experimental structure or 3D structure model of the protein is known, this can be entered via a Protein Databank (PDB) identifier [20]. In the EVFold mode, the server assesses the accuracy of structure prediction from sequence alone by comparing the EVfold predicted structure to the known structure using standard 3D superimposition methods, such as the one in the PyMOL molecular graphics software. In the EVcouplings mode, the server can map ECs onto the known structure for functional interpretation, without 3D structure prediction.

### 2.3 Algorithm: maximum-entropy model for protein sequences

The algorithm uses a by now well-established [3,7,21–24] maximum-entropy probability model to identify ECs in the protein family. It uses a subset of the ECs to impose residue pair distance constraints in the molecular modelling software CNS [25] to generate 3D structures using distance geometry projection followed by (moderately long, as of March 2015) simulated annealing by molecular dynamics.

The server has a choice of two levels of approximation for inferring the parameters of the global, maximum-entropy probability model: “DI”, a mean-field based coupling analysis with parameters obtained by inversion of the covariance matrix and direct information scoring using an analogue of mutual information with direct information marginal probabilities [7, 21]; and “PLM”, a pseudo-likelihood maximization approximation with corrected norm scoring [23]. DI, also called mean field DCA, tends to be the faster but less accurate method compared to PLM. For a review of methods see [24]. We recommend the slower but more accurate PLM as default.

### 2.4 Output: evolutionary couplings and/or predicted 3D structures

The server provides a list of evolutionarily coupled residue pairs ranked by EC score, 2D maps of predicted contacts and/or a set of predicted all-atom 3D structures. High-ranking ECs represent strong evolutionary constraints and tend to reflect residue proximity in the folded protein structure and/or functionally important interactions for any protein in the family. The ECs are visualized as a 2D contact map and also mapped onto a known or predicted 3D structure of the target protein, for example as lines connecting two residues. The total strength of ECs affecting a single particular residue is visualized as a residue property in 3D, with the aggregate EC strength (summed over all partner residues) as residue atomic sphere color (Figure 2) or residue thickness in “sausage” mode.

**Figure 2:**
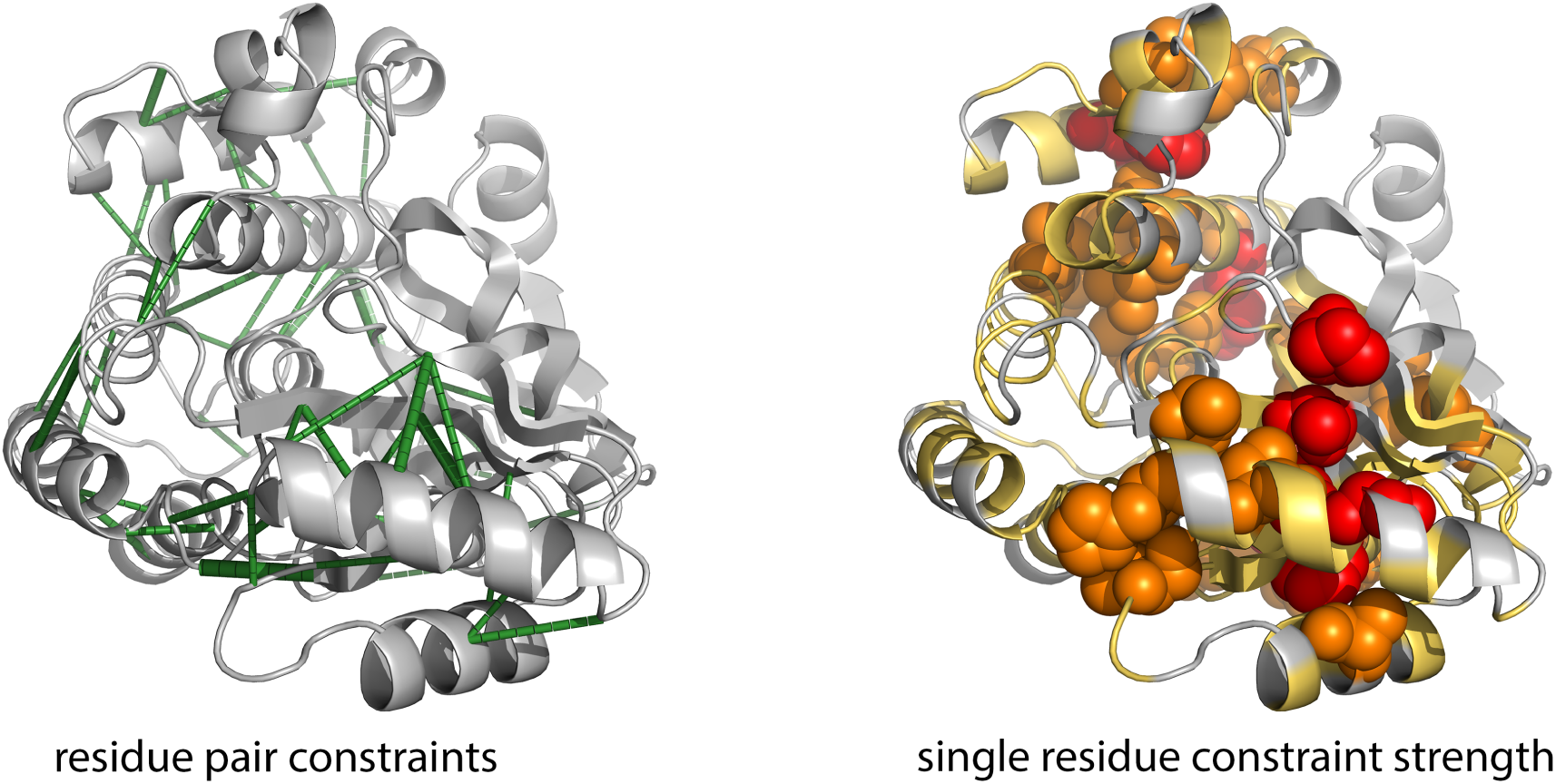
Evolutionary couplings visualized on known structures or predicted structures are informative about essential residue-residue interactions (option EVcouplings for known structures, EVfold for unknown structures). Many strong residue pair couplings (left, green lines) in a bacterial protein transacylase (Uniprot: FABD_ECOLI; PDB: 1MLA) link residues that are in contact in 3D. Residues with strong single residue constraint strength (ECs summed over all coupled residue partners) may point to interaction sites that implement evolutionary requirements (the most strongly coupled residues as red spheres, followed by orange spheres for medium strength, and yellow ribbon for low strength).

To reflect modelling uncertainty for a given set of distance constraints and stochastic decisions in the simulated annealing protocol, the server typically computes several hundred models for a given target protein, e.g, ∼2L models for a protein of length L. The models are ranked by a scoring function that estimates the likelihood of being a correct and typical protein structure, using criteria based on distributions of dihedral angles, torsion angles, solvent accessibility, constraint satisfaction, and agreement with predicted secondary structure. The model scoring function is under development and improvements are expected in a future version of the server.

### 2.5 Current status of prediction accuracy

For a blinded benchmark set (testing 3D prediction on proteins of known 3D structure without using any information from the known structure) good 3D models (TM-score ≥0.5 [28] can be predicted, in the current implementation, in about half of all cases (Table 1). For this test, we used a structurally representative set of domains of known structure (using the CATH database as a guide [29]) and, following sequence searches and MSA construction, used a stringent cutoff on the minimal depth and breadth of the multiple sequence alignment. In particular, we required that the effective number of non-redundant sequences (Meff) normalized by protein length (L) is greater than 4.0 (depth) and that the coverage of the protein domain being modeled by non-gappy alignment columns exceeds 75 percent (breadth). In the reduced set of 63 protein domains (out of 140), which exceeded these thresholds, 38 (60%) had very good prediction accuracy (TM-score >= 0.5), while 25 (40%) had accuracy below the customary threshold (TM-score < 0.5). Deeper alignments (more sequences available in the family) tend to lead to better prediction accuracy. As a rule of thumb, proteins above these thresholds yield good predictions in about half the cases (Figure 3, Table 1).

**Figure 3:**
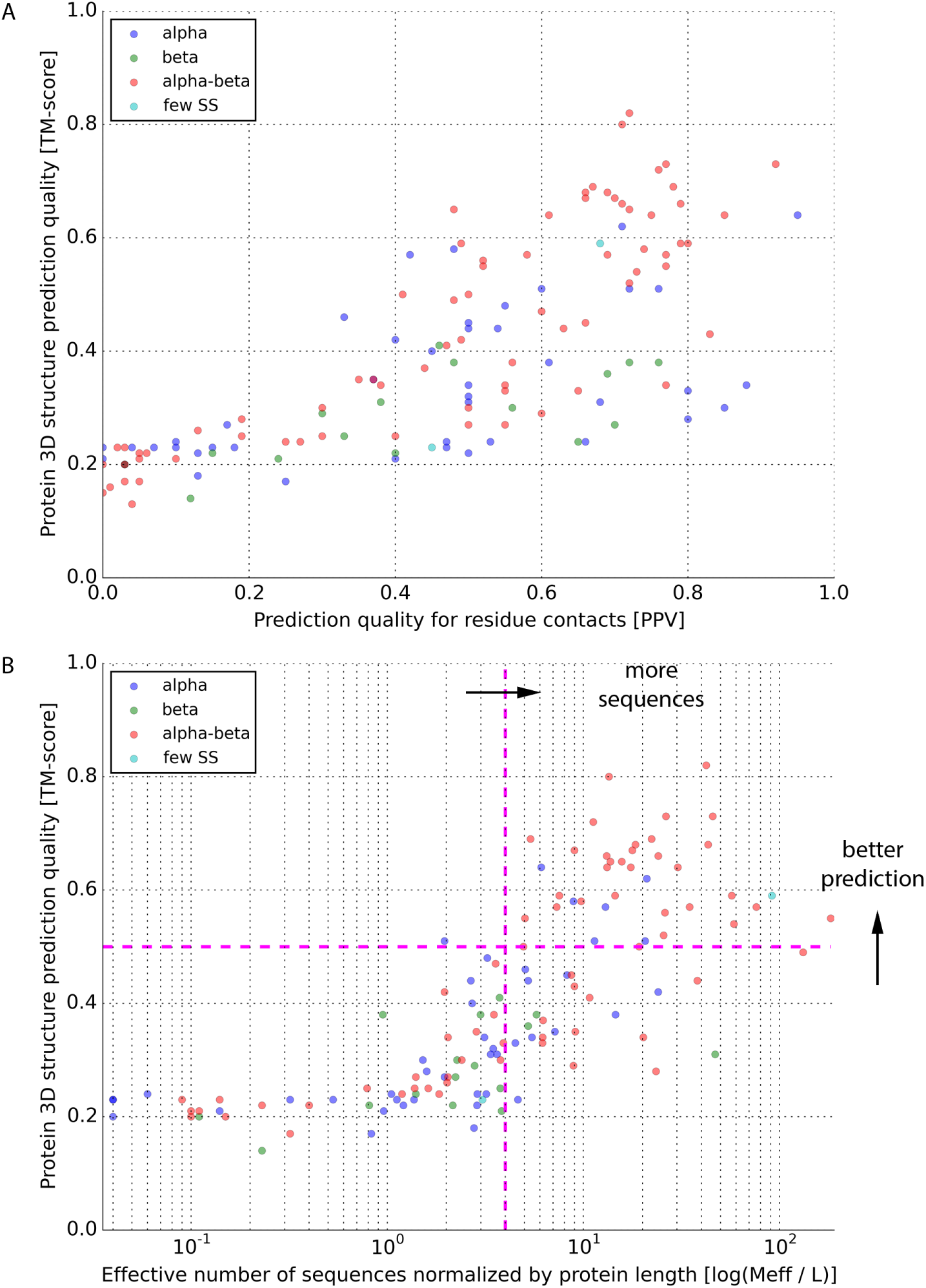
EVfold benchmark on 140 proteins of known structure indicates requirements for the amount of sequence information for good predictions. Folding results from proteins (dots) with diverse CATH topologies. (A) 3D structure prediction quality as measured by TM-score as a function of the prediction quality for residue contacts as measured by PPV. (B) 3D structure prediction quality as measured by TM-score as a function of the effective number of sequences normalized by protein length. Proteins in the upper right quadrant (‘well folded’ in Figures 4-6) were well predicted. Proteins in the lower right quadrant were less well predicted (‘less well or badly folded’ in Figures 7-9) although they passed the threshold for sufficient sequence information. Runs were done at E-value 10^−4^, 5 jackhmmer iterations, and filtering of alignment columns and rows if gap content exceeded 30 percent. PPV is the positive predictive value (number of true positives divided by the sum of true and false positives). TM-score is the template modeling score (>0.5 considered good). Results are for the best model within top 10 ranked. Secondary structure protein fold types are alpha helical (blue), beta-strand (green), alpha-beta mix (red), few secondary structures (‘few SS’, teal).

**Table 1.**
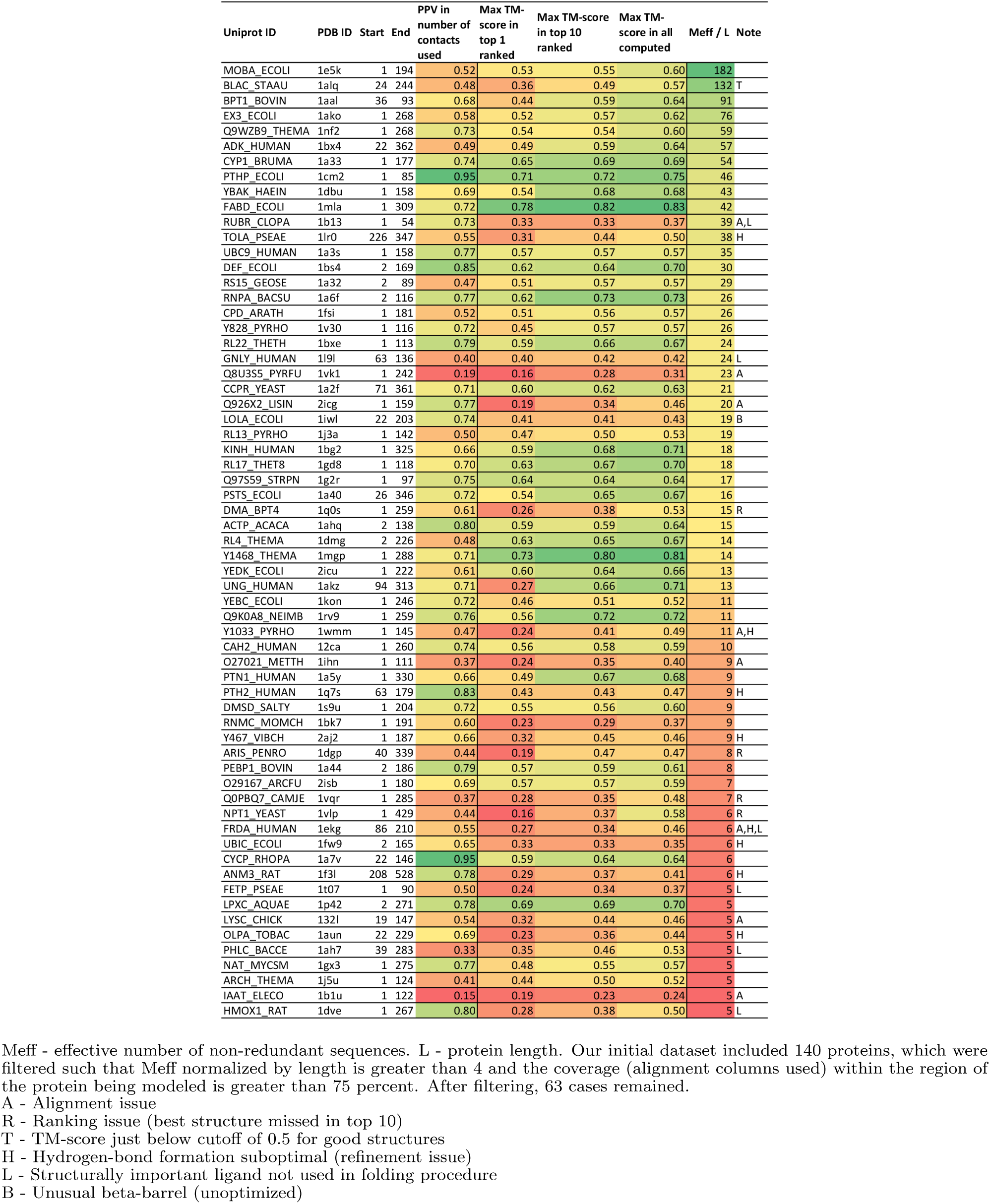
Performance of EVfold on a diverse set of proteins.

While the TM-score as a quantitative measure of 3D structure prediction accuracy is useful, and often used, it does not fully reflect the essence of the success of the EVfold method, which in many cases correctly predicts the topographical arrangement, i.e., the relative spatial arrangement of secondary structure elements. In some cases predicted structures with TM-score < 0.5 have perfect prediction of topography, while for TM-score > 0.5 there can be topographical errors. We therefore provide, for intuitive inspection by the reader, a number of explicit examples of predicted structures in the ‘well folded’ (Figures 4-6) and ‘less well or badly folded’ (Figures 7-9) categories, both as 2D contact maps as well as 3D structural superimpositions on the known structure. Coordinates for these structures are on the EVfold.org web site. While a purist comparison of prediction accuracy would only assess the top-ranked predicted structure, we typically compute prediction error in the blinded test for the actually best structure in the top ten ranked ones, on the grounds that serious exploration of predicted structure, for functional interpretation for example, can afford scanning through ten predicted structures. Prediction accuracy for the single top-ranked structure are provided for completeness.

**Figure 4:**
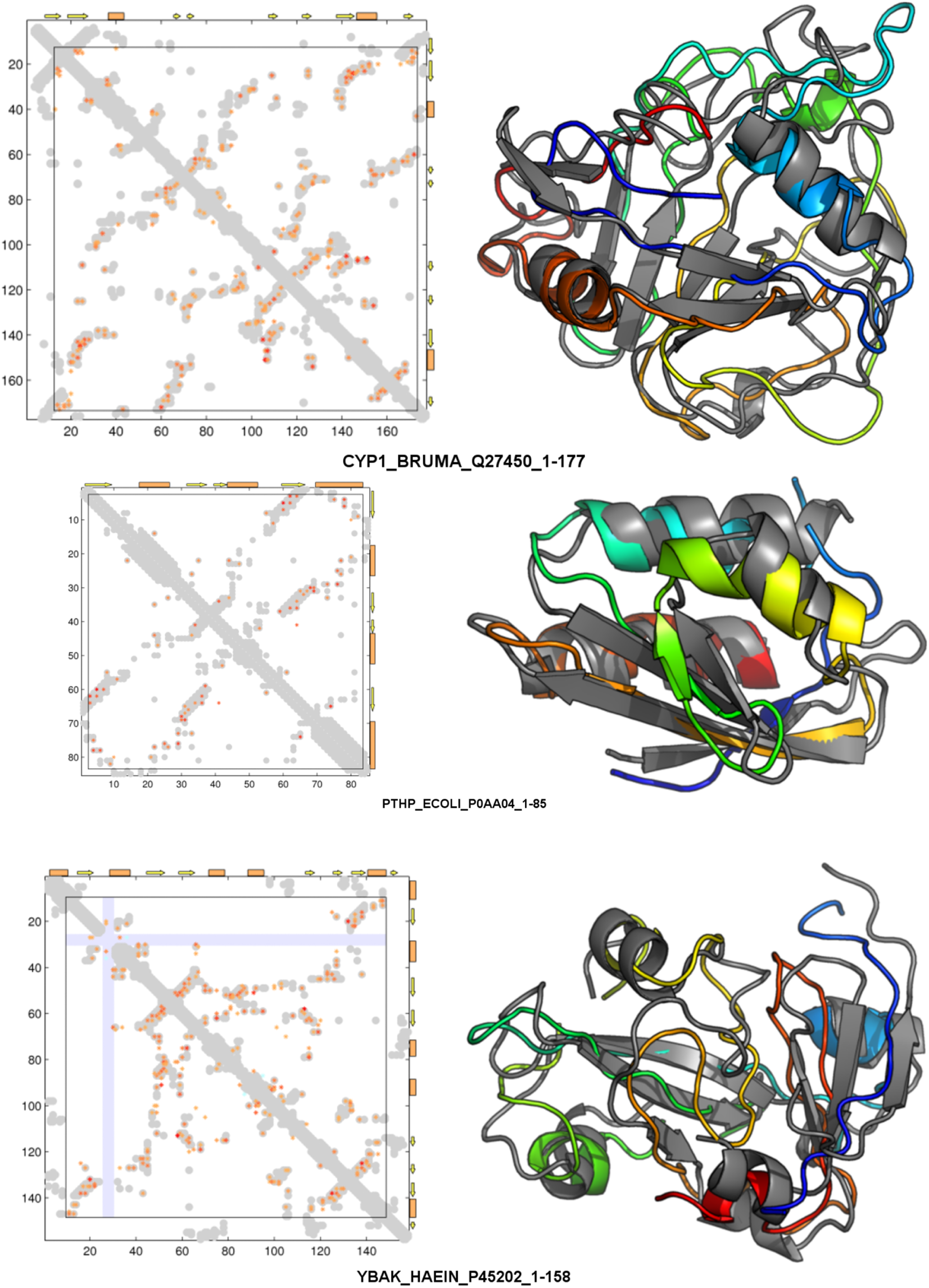
Well folded proteins. A selected subset of proteins in Table 1 with TM-score above 0.5 (best predicted by TM-score, sorted by Meff/L). Protein names (example): Uniprot Name_Species (CYP1_BRUMA), Uniprot ID (Q27450), residue range from-to (1-177). Left: Quality of contact prediction is higher the more predicted contacts (red to orange as EC value decreases) match the contacts derived from the experimental structure (grey). No experimental information is available in segments of the protein missing in the reference (PDB) structure (light blue ribbons). Contact patterns parallel or antiparallel to the diagonal are contacts between secondary structure helices (orange rectangles) or beta strands (arrows). Right: predicted 3D structure (rainbow-colored cartoon) superimposed against reference experimental structure (grey ribbon).

**Figure 5:**
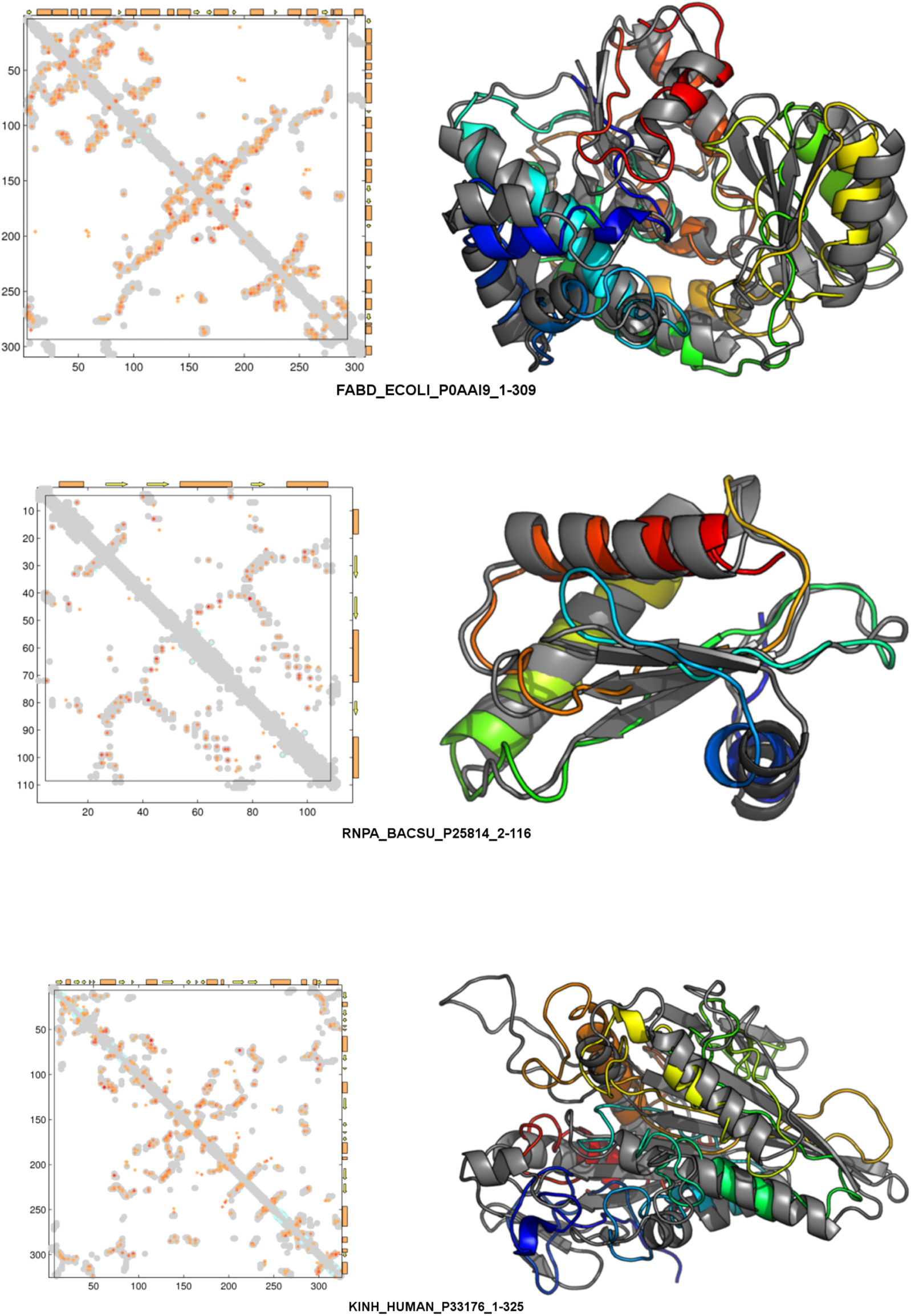
Well folded proteins (cont’d). Coloring as above.

**Figure 6:**
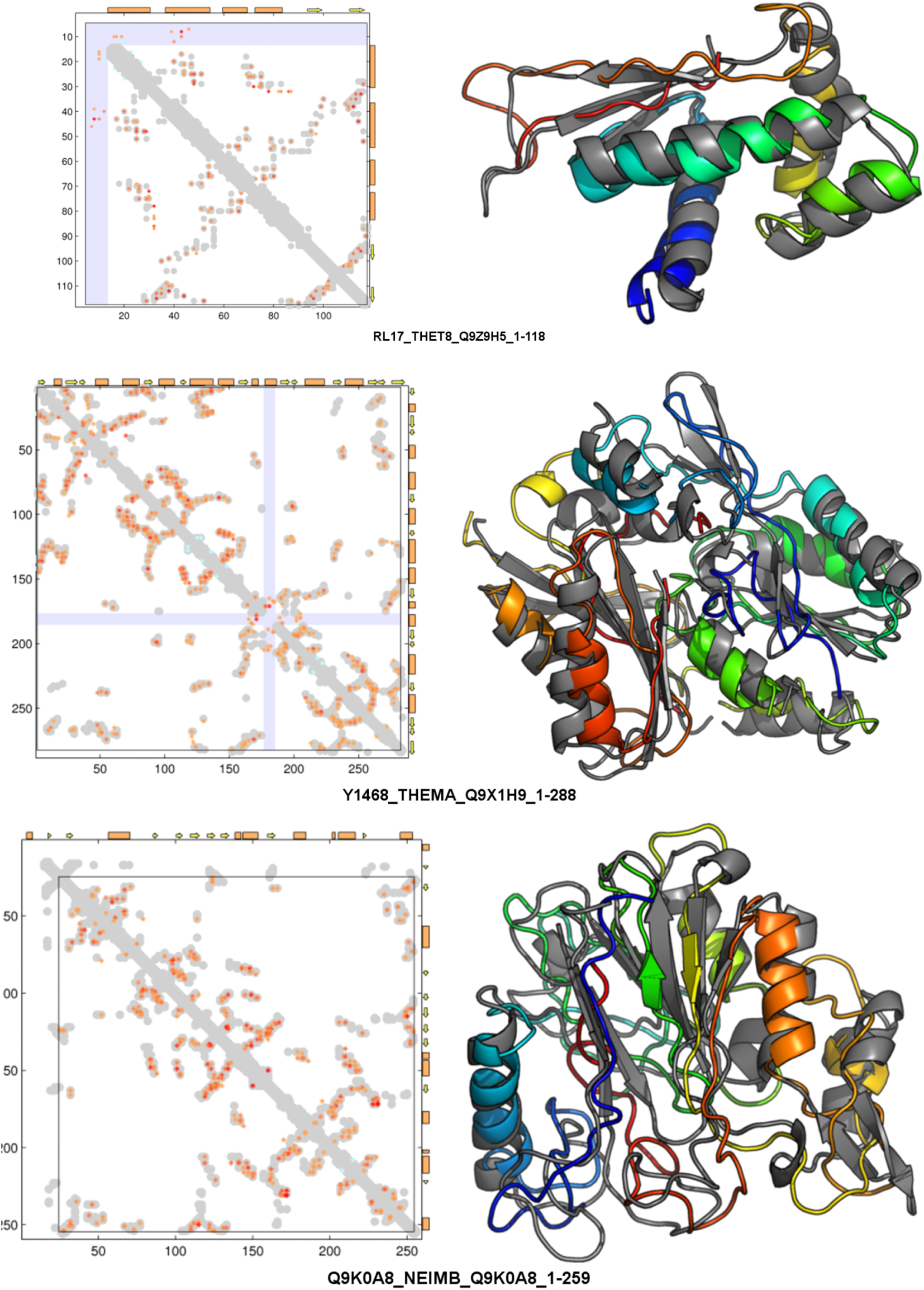
Well folded proteins (cont’d). Coloring as above.

**Figure 7:**
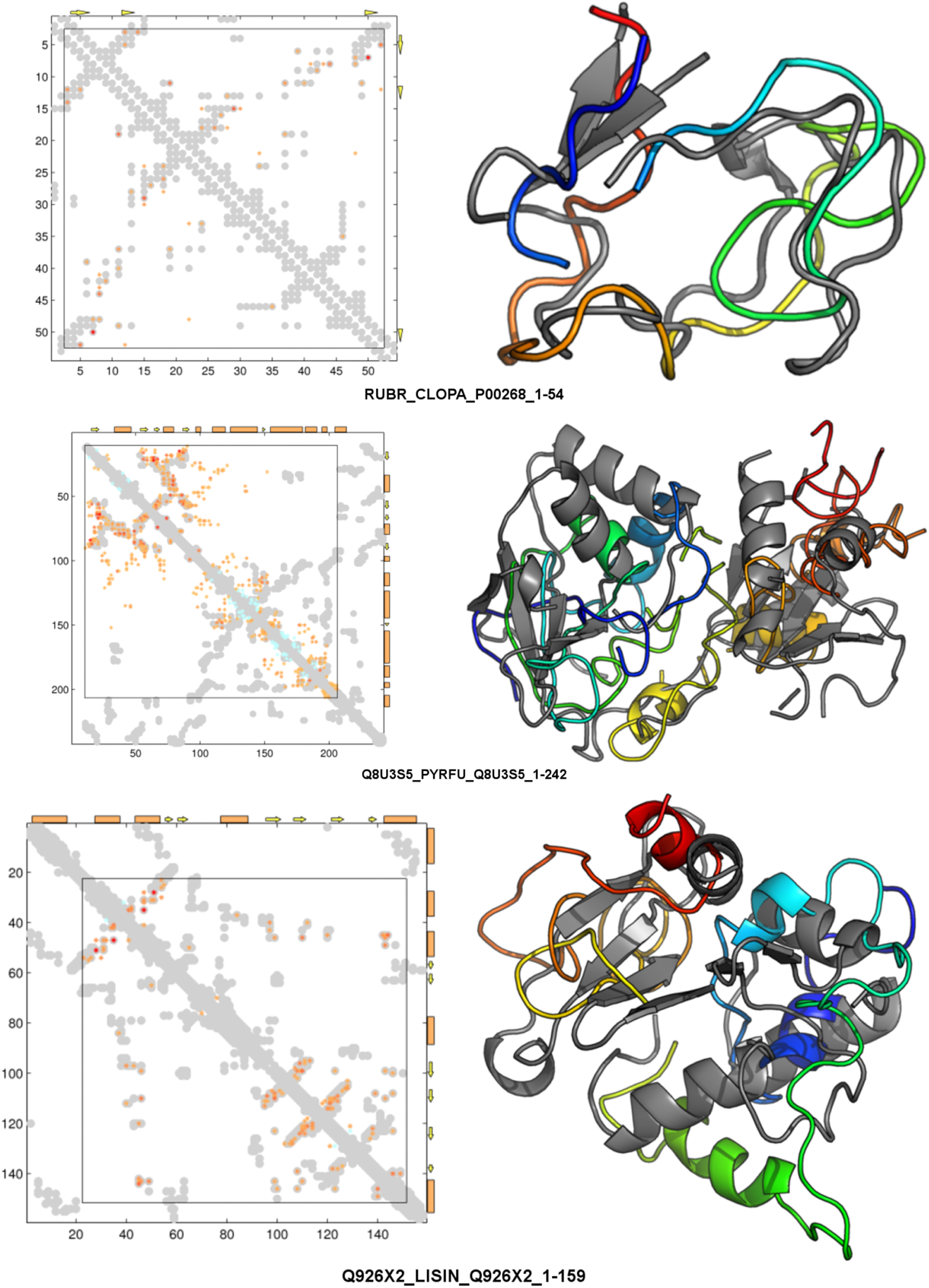
Less well or badly folded proteins. A selected subset of proteins in Table 1 with TM-score below 0.5 (least well predicted by TM-score, sorted by Meff/L). Graphical representation as in previous figure.

**Figure 8:**
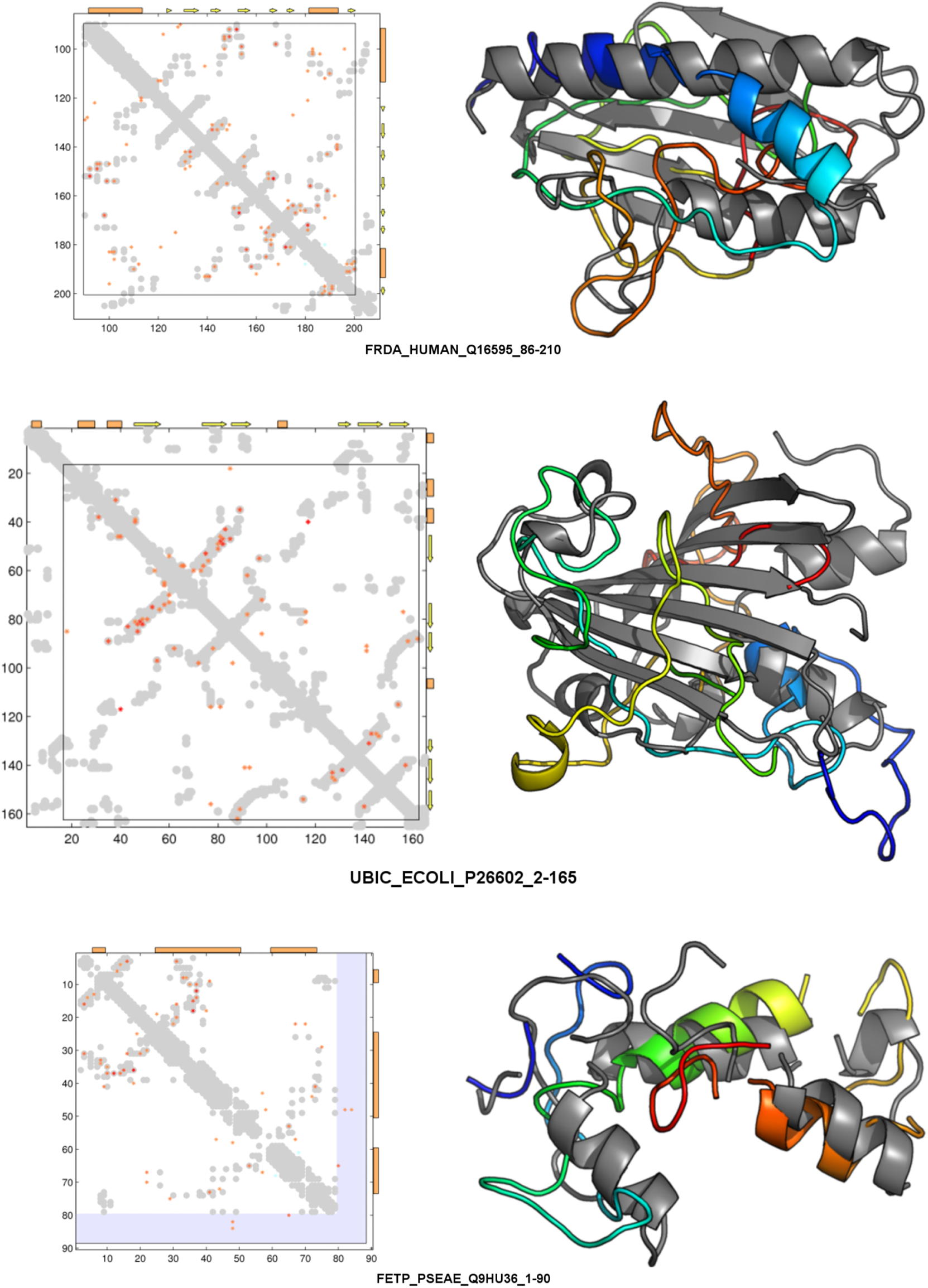
Less well or badly folded proteins (cont’d). Coloring as above.

**Figure 9:**
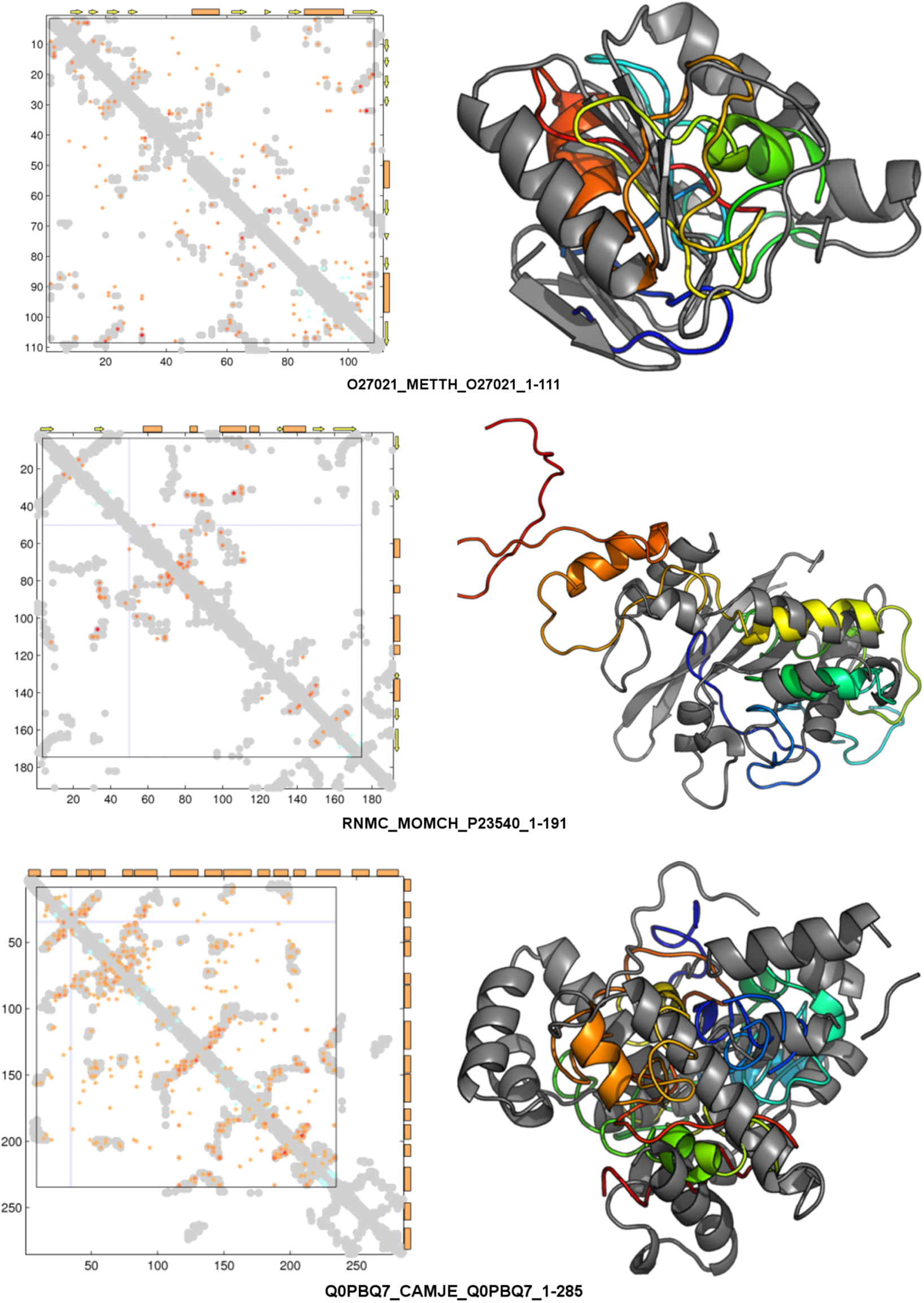
Less well or badly folded proteins (cont’d). Coloring as above.

### 2.6 Factors affecting prediction accuracy and future improvements

The most important factor affecting 3D structure prediction accuracy is the depth and diversity of sequence information in the family containing the target protein domain. Inspection of about 50 cases, anecdotally, suggests additional factors related to lower prediction success, such as unusual amino acid composition or disulfide patterns, apparently erratic sections of the MSA, or apparently discontinuous subfamily structure. In some cases, predicted structures in very good agreement with experimental structures were at a low rank in the set of all computed structures (generally ∼2L), indicating less than fully adequate criteria in the ranking score. In some cases of correctly predicted topography of structure segments, well-placed beta-strands did not have the expected pattern of well-formed hydrogen bonds. Therefore, three important areas of desired improvement are: improved alignment procedure (cutoffs, filters, uneven sequence weights); improved refinement of the 3D coordinates of the protein models using more elaborate simulated annealing by molecular dynamics; and improved ranking criteria for structures in the set of predicted structures for a target protein.

## 3 RECOMMENDATIONS

Before submitting a protein for 3D structure prediction, it is useful to first check for protein domains in the target sequence, via the Pfam database for example, and consider submitting each domain separately. For each domain, one should check whether a 3D structure or that of a homologous ‘template’ is available, which would directly lead to a 3D structure model. Such models are often already deposited in the very useful database of protein models, www.proteinmodelportal.org, or can be generated using existing tools such as HHpred [30]. For any prediction or EC analysis run, one can check the alignment quality by inspection, as this is the single most important factor affecting predicition quality. We suggest a minimum number of ∼5L diverse sequences (depth) and coverage of the target domain over at least ∼0.75L (breadth). Once a contact map (top ECs) is generated, visual inspection can provide useful clues as to likely prediction success, looking for ‘protein-like’ structured patterns in the 2D predicted contact map. Caution is needed for homo-multimers and alternative conformations, as inferred ECs from the target sequence may reflect contacts between monomers of a homo-oligomer or contacts in alternative conformations, such as closed and open forms of a channel. The server returns a ranked set of 3D models for the target protein, typically several hundred; it is reasonable to inspect the top ranked model and perhaps about a dozen top additional ones. In our view, both the ranking score as well as the 3D structure refinement process can be substantially improved (work in progress). In summary, currently about one in two proteins with alignment quality above the recommended thresholds lead to excellent predictions. The number of proteins accessible to the method is rising rapidly as genome sequencing accelerates.

## 4 ACKNOWLEDGEMENTS

### Author contributions

RS and LY developed the backend; YS developed the front end; YA, RS developed MSA viewer; TH, DSM developed the transmembrane pipeline; RJF, CS, SH collected and analyzed the protein datasets; RJF, CS, RS wrote the text; all authors edited the text; CS, DSM conceived of and designed the project.

## Funding Support

We thank Richard Stein for helpful discussions and feedback. This work was funded by the US National Institutes of Health (R01GM106303 to CS at MSKCC and DSM at HMS) and the Canadian Natural Sciences and Engineering Research Council (fellowship to RJF).

